# Exposure to temporal randomness promotes subsequent adaptation to new temporal regularities

**DOI:** 10.1101/2023.06.04.543595

**Authors:** Orit Shdeour, Noam Tal-Perry, Moshe Glickman, Shlomit Yuval-Greenberg

## Abstract

Noise is intuitively thought to interfere with perceptual learning; However, human and machine learning studies suggest that, in certain contexts, variability may reduce overfitting and improve generalizability. Whereas previous studies have examined the effects of variability in learned stimuli or tasks, it is hitherto unknown what are the effects of variability in the temporal environment. Here, we examined this question in two groups of adult participants (N=40) presented with visual targets at either random or fixed temporal routines and then tested on the same type of targets at a new nearly-fixed temporal routine. Findings reveal that participants of the random group performed better and adapted quicker following a change in the timing routine, relative to participants of the fixed group. Corroborated with eye-tracking and computational modeling, these findings suggest that prior exposure to temporal randomness promotes the formation of new temporal expectations and enhances generalizability in a dynamic environment. We conclude that noise plays an important role in promotion perceptual learning in the temporal domain: rather than interfering with the formation of temporal expectations, noise enhances them. This counterintuitive effect is hypothesized to be achieved through eliminating overfitting and promoting generalizability.

## Introduction

Sitting in the driver-seat waiting for the streetlight to change, holding up a hand waiting for a teacher to call on us, or standing at the bus-stop waiting for the bus to arrive; our daily lives are full of scenarios in which expectations shape our behavior (Nobre et al., 2007). The formation of new expectations depends on perceptual learning, as it requires prior experience with specific statistical rules (Nobre & Van Ede, 2018). Decades of research on perceptual learning examined how experience with stimuli and tasks shapes perception, attention, and behavior (Fahle et al., 2002; Goldstone, 2003; Sagi, 2011; Sasaki et al., 2009; Sigman & Gilbert, 2000). These studies have consistently shown that, following practice, there is a typical sustained improvement in performance on perceptual tasks. However, this improvement is often very specific – performance improves on the trained task with the trained set of stimuli, but rarely on other, even similar, tasks or stimuli (Maniglia & Seitz, 2018; Sagi, 2011; Wright & Zhang, 2009). This consistent finding raises an important question – if constructing expectations requires learning from prior experience and if such learning is so specific, how does the cognitive system construct reliable expectations in a dynamic environment in which statistical rules often change?

This intriguing question occupies not only cognitive scientists but also computer scientists. A widely discussed topic in the field of machine learning is the tradeoff between overfitting and underfitting – on the one hand, the goal to optimally adjust the learning models to the trained exemplars, but on the other hand, the risk that these models will over-adjust to old exemplars, thereby hindering their ability to generalize to new ones (Wang & Zhong, 2003). Such overfitting is the consequence of learning statistical rules that may apply to the sub-set of trained exemplars but are irrelevant to subsequent new exemplars, resulting in poor predictions. One mean by which the overfitting risk can be eliminated is by adding random noise to the trained exemplars, thereby reducing the fit of the algorithms to the trained data while improving their generalizability (Bishop, 1995a, 1995b; James et al., 2013).

Adding noise to learning processes is, intuitively, expected to disrupt learning or at least slow it down. However, as we know that adding variability enhances the generalizability of learning algorithms, it could be hypothesized to have a similar, positive, effect on human learning. Existing evidence suggests that when a few skills, perceptual tasks, or features are interleaved during practice, the level of learning and its generalizability to untrained items are improved (Brady, 1998; Harris et al., 2012; Magill & Hall, 1990; Szpiro et al., 2014; Wang et al., 2012; Wright et al., 2010; Xiao et al., 2008). However, it is still an open question whether adding temporal randomness to the environment could be advantageous for learning. Specifically, it is unknown whether temporal noise would improve adaptation to a new temporal regularity. While previous studies have stressed the importance of the particularities of training conditions to generalizability and overfitting of learning (e.g., Hung & Seitz, 2014), the crucial role of the temporal environment in which learning takes place has so far been generally overlooked.

The goal of this pre-registered study was to examine this question focusing on temporal expectation. To this end, we used the *foreperiod paradigm* which creates an association between a warning signal (cue) and a target, with a fixed or varying interval between them called a *foreperiod* (Baumeister & Joubert, 1969; Coull & Nobre, 1998; Näätänen, 1970; Niemi & Näätänen, 1981). Here, we employed this paradigm with two groups of participants who performed a visual discrimination task. In the first phase of the experiment (*acquisition phase*), participants of one group (*the random group)* were exposed to random foreperiods (1700-3700 ms), while participants of the second group (*the fixed group*), were exposed to fixed foreperiods of 2700 ms. In the second phase (*transfer phase*), both groups performed the same task with a nearly-regular temporal foreperiod regime of 80% 700 ms and 20% 2700 ms (see **Fig. 1A**).

**Fig 1.**
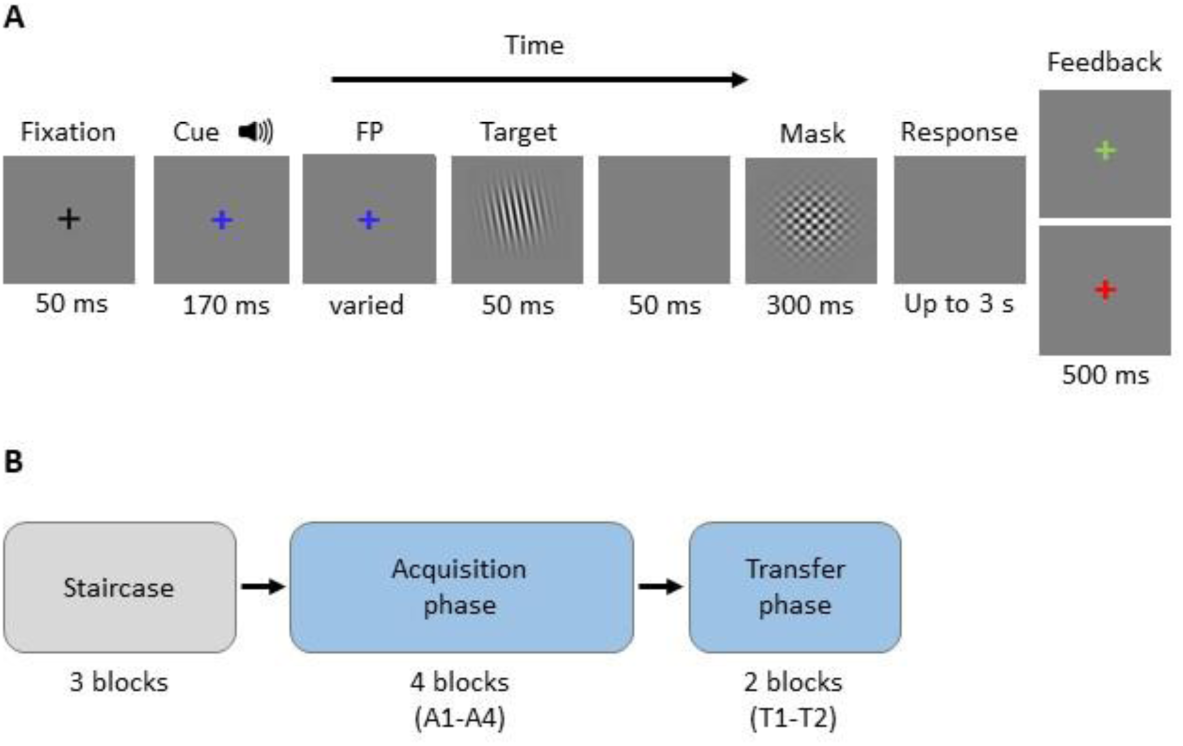
Procedure. (**A**) Trial procedure. Every trial began with a black fixation cross followed by a simultaneous presentation of auditory (pure tone) and visual (blue fixation cross) cues. The blue fixation cross continued to be presented during the interval between cue onset and the target onset (foreperiod, FP). The duration of the foreperiod varied according to the phase and group conditions. After the foreperiod, the target was presented and was followed by a mask. Following target presentation, participants were asked to determine whether the target was tilted left or right and respond by pressing one of two buttons, and then received feedback (green or red fixation cross for correct or incorrect responses, respectively). After the feedback, there was a variable interval of 0.2-0.7 s before the next trial was initiated. **(B)** Experimental blocks. The experiment included a total of nine blocks: In three preliminary staircase blocks, the tilt threshold was chosen individually per participant to achieve approximately 80% accuracy rates. This was followed by the main experiment that included four acquisition blocks, and two transfer blocks. Each experimental block included 96 trials.

The behavioral findings of this study, corroborated by eye-tracking and computational modeling, revealed that participants of the random group showed a higher ability to adapt to a new temporal routine, relative to those of the fixed group. This confirms the hypothesis that in human learning, as in machine learning, prior exposure to a noisy environment reduces overfitting and improves the ability to adapt to new statistical rules. This finding highlights the beneficial role of noise in the adaptation of predictions and expectations according to a changing environment. The realization that adding noise is not always harmful and could even be beneficial to behavior, has wide implications for developing noise-based education and rehabilitation procedures.

## Results

### Accuracy rates

Accuracy results were analyzed by constructing a three-way GLM model assuming a binomial family of response and a logit link. The model includes the fixed factors Trial Number (modeled with 1st polynomial, i.e., linear component), Group (fixed/ random) Phase (acquisition/ transfer) and their accompanying interaction terms, allowing for full random effect structure (random intercept, random slopes for each variable and their interactions, and accompanying correlation parameters). For the main analysis we included all the trials of the acquisition phase and only trials of foreperiod 700 ms of the transfer phase. In a complementary analysis (see below) we have separately examined transfer trials of 2700 ms.

This GLMM analyses revealed no main effect for Group (χ^2^(1) = 0.311, *p* = .577) nor for Trial number (χ^2^(1) = 0.813, *p* = .367), and the two factors did not significantly interact with each other (χ^2^(1) = 0.021, *p* = .886). However, there was a significant main effect for Phase (χ^2^(1) = 25.267, *p* < .0001), with an overall decrease in accuracy at the transfer phase (M=70.8%, SD = 12.0%) relative to the acquisition phase (M=80.4%, SD = 7.2%), indicating that the change in the foreperiod distribution from the acquisition to the transfer phase interfered with performance. The phase effect was significantly modulated by Trial number (χ^2^(1) = 7.755, *p* = .005). This interaction resulted from a shallow decrease of accuracy over trials during the acquisition phase (log estimate = –.077, Wald’s z = –2.049, *p* = .040), possibly due to fatigue, and an overall marginal increase in accuracy during the transfer phase (log estimate = .104, Wald’s z = 1.835, *p* = .067), possibly due to the gradual adaptation for the new foreperiod routine.

Critical for the purpose of this study is the significant two-way interaction between Group and Phase (χ^2^(1) = 8.291, *p* = .004). A follow-up analysis performed separately on the two phases revealed that accuracy rates for the two groups were similar in the acquisition phase (log estimate = –.105, Wald’s z = –.747, p = .455), but differed significantly in the transfer phase (log estimate = –.483, Wald’s *z* = –2.559, *p* = .010). The random group showed higher accuracy in the transfer phase (M=74.9%, SD = 12.7%) than the fixed group (M=66.9%, SD = 10.1%; **Fig 2**), consistent with our hypothesis that exposure to random intervals improves adaptation relative to fixed intervals.

**Fig 2.**
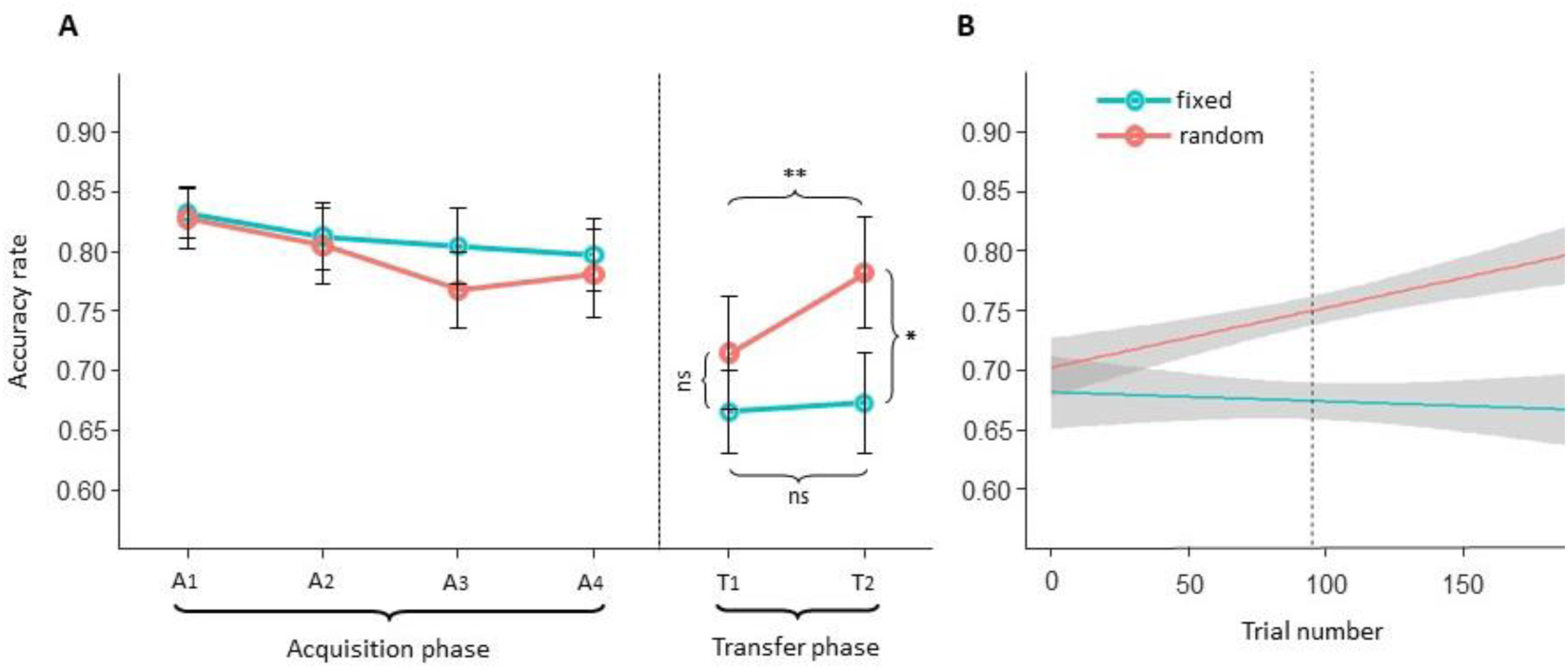
Accuracy rate results. (**A**) The X-axis represents the experimental blocks in chronological order (Acquisition= A 1-4, Transfer= T 1-2). Dashed line represents the onset of the transfer phase. Error bars depict ±1 standard error from the group mean. Simple effects are marked as: p<0.05*, p<0.01**, ns=not significant. **(B)** Regression line of trial-wise temporal dynamic for accuracy rates in each group during the transfer phase, based on the individual participant data across trials. Shaded area represents 95% confidence interval around the slope. Dashed line represents the onset of second block (T2).

Furthermore, we observed a significant three-way interaction between Phase, Group, and Trial number (χ^2^(1) = 4.546, *p* = .033), indicating that the modulation of Group by Phase evolved over trials. Again, we performed a follow-up analysis separately on the two phases. In the acquisition phase, both groups showed a similar pattern of performance over trials, including a gradual decrease in performance (random group: log estimate = –.055, Wald’s z = –1.041, *p*= .298; fixed group: log estimate = –.099, Wald’s *z*= –1.856, *p* = .063), which was likely due to fatigue. In contrast, during the transfer phase (see **Fig. 2B**), the groups differed in their performance dynamics over trials. Whereas at the beginning of the transfer phase, both groups showed a sharp decline in performance relative to the acquisition phase (see **Fig. 2A**); the fixed group remained at a low and stable performance level throughout the phase (log estimate = 0.002, Wald’s *z* = 0.003, *p* = .976) while the random group quickly recovered and showed an overall gradual enhancement over trials (log estimate = 0.205, Wald’s *z* =2.489, *p* = .013). These findings support the hypothesis that learning a systematic interval routine hinders participants’ ability to adapt to new routines.

In a complementary analysis, we examined the minority (20%) of transfer phase trials with a foreperiod of 2700 ms. We constructed the same GLM model as in the main analysis and included all trials of the acquisition phase and only the 2700 ms trials of the transfer phase. Unlike the previous model, the present model includes a random structure of random intercept and random slopes of RT, Trial number and Phase, as well as the interaction between trial number and phase.

As in the main analysis, we found no evidence for a Group main effect (χ^2^(1) = 0.069, *p* = .793), and for an interaction between Group and Trial number (χ^2^(1) = 0.058, *p* = .810). Also, similarly to the previous analysis, we found a main effect of Phase (χ^2^(1) = 11.357, *p* < .0001), caused by an overall decrease in accuracy at the transfer phase compared to the acquisition phase, and an interaction of Phase and Trial Number (χ^2^(1) = 10.801, *p* = .001). This interaction resulted from a decrease in accuracy over trials during the acquisition phase (log estimate = –.078, Wald’s z = – 2.136, *p* = .032) and no evidence for such a decrease during the transfer phase (log estimate = –.100, Wald’s z = 1.428, *p* = .153). We also found a significant interaction between Group and Phase (χ^2^(1) = 6.088, *p* = .001). As in the main analysis, both groups performed the task similarly during the acquisition phase (log estimate = .107, Wald’s z = 0.759, p = .448), but unlike the main analysis, we found only a marginally significant difference between the groups in the transfer phase (log estimate = –.344, Wald’s z = –1.703, p = .089). Unlike the main analysis here we found a main effect of Trial number (χ^2^(1) = 16.771, *p* < .001), caused by a reduction in accuracy over trials. The three-way interaction of Phase, Group, and Trial number was found to be only marginally significant (χ^2^(1) = 3.678, *p* = .055), but we have nevertheless performed follow-up analysis examination of performance of the two groups in the transfer phase. This analysis revealed that, similarly to the main analysis, the performance of the fixed group remained unchanged over trials (log estimate=-0.006, Wald’s *z*=-0.062, *p*= .095) while the random group marginally increased over trials (log estimate = 0.206, Wald’s *z* =1.991, *p* = .046). This is an important finding because the foreperiod of 2700 ms was the foreperiod on which the fixed group trained during the acquisition phase. These findings suggest that the fixed group did not benefit relative to the random group, even in trials that maintained the temporal routine that they were initially exposed to.

### Speed-accuracy tradeoff

In this study we used a difficult, non-speeded, task that is more suitable for studying accuracy rates and less for studying RTs. We, nevertheless, examined RTs to ensure that the observed accuracy-rate findings cannot be explained solely as speed-accuracy tradeoffs (Carrasco et al., 2004; Giordano et al., 2009; Wickelgren, 1977). To this end, we inserted the RT results as an additional fixed factor in the accuracy model described above, along with its interactions with the other factors.

We found a main effect of RT on accuracy resulting from a decrease in RT when accuracy was higher (χ^2^(1)=53.072, *p*<.0001). This is opposite the effect that could have resulted from a speed-accuracy trade-off, suggesting that no such trade-off exists in these data. Furthermore, the main finding of this research – a significant three-way interaction of Phase X Group X Trial number – was found with even this new model which includes RT (χ^2^(1)=4.008, *p*= .045), suggesting that this effect is independent of RT. A follow-up examination, performed separately on accuracy rates of the two phases and groups revealed opposite effects of RT on accuracy relative to the ones expected by speed-accuracy tradeoff, i.e. accuracy was lower when RT was higher. This was found for both groups and phases (fixed group, acquisition phase: log estimate=-0.540, Wald’s *z*=7.453, *p*<.001, fixed group transfer phase: log estimate=0.393, Wald’s *z*=-5.225, *p*< .001, random group, acquisition phase: log estimate=-0.634, Wald’s *z*=-8.749, *p*<.001, random group, transfer phase: log estimate=-0.400, Wald’s *z*=-4.991, *p*<.001). These findings are consistent with our main findings, suggesting that they are independent of RT and cannot be explained by a speed-accuracy tradeoff. This conclusion is also supported by the result of a drift-diffusion model (DDM, described below), which takes both RT and accuracy rates into account. Following our pre-registered plan, we have also performed a full GLM of RT, which is provided in Supplementary materials S2. The RT results during the transfer phase are presented in **Fig 3**.

**Fig 3.**
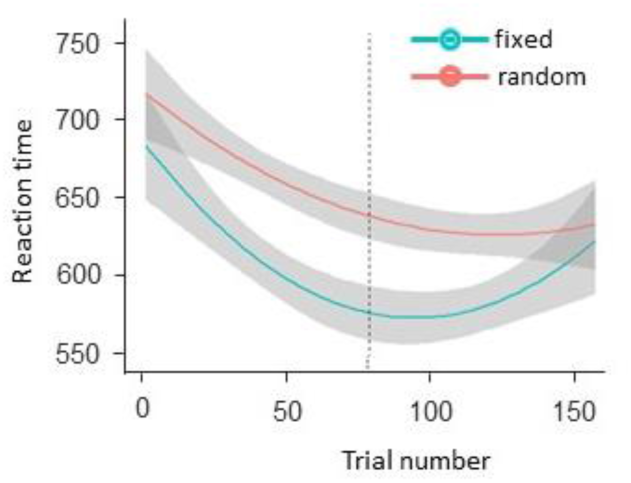
Reaction time results. Regression line of trial-wise temporal dynamic of reaction times in the two group (Fixed and Random) during the transfer phase, based on individual participant data across trials. Shaded area represents 95% confidence interval around the slope. Dashed line represents the onset of the second block (T2).

### Eye movements

In a series of previous studies, we showed that the mean saccade-rate at –100 to 0 relative to target onset (pre-target SR), can be used as an index for temporal expectation: pre-target SR was consistently found to be lower for expected relative to unexpected targets (Amit et al., 2019; Dankner et al., 2017; Tal-Perry & Yuval-Greenberg, 2020, 2021). Here, we used this measurement to assess temporal expectations across the phases and in the different groups. We constructed a three-way GLM model with the dependent variable of pre-target SR (mean saccade rate at 0-100 ms relative to target onset), assuming a binomial family of response and a logit link. The model includes the fixed factors Trial Number (modeled with 1st polynomial, i.e., linear component), Group (fixed/ random) Phase (acquisition/ transfer), and their accompanying interaction terms, allowing the random effects of the random intercept and the random slopes of Trial number and Phase. Similar findings were obtained using an ANOVA, as in the pre-registered plan (Supplementary material S2).

We found a main effect of Phase (χ^2^(1) = 8.211, *p* = .004). There was no significant main effect of Group (χ^2^(1) = 0.238, *p* =.625) nor of Trial number (χ^2^(1) = 1.141, *p* =.285), but there was a significant interaction between Phase and Group (χ^2^(1) = 24.413, *p* <.0001), indicating that mean pre-target SR of the random group decreased in the transfer phase relative to the acquisition phase, but increased in the fixed group (**Fig 4**). There was no interaction between Trial number and Group (χ^2^(1) = 1.325, *p* =.250) nor between Trial number and Phase (χ^2^(1) = 0.243, *p* =.622).

**Fig 4.**
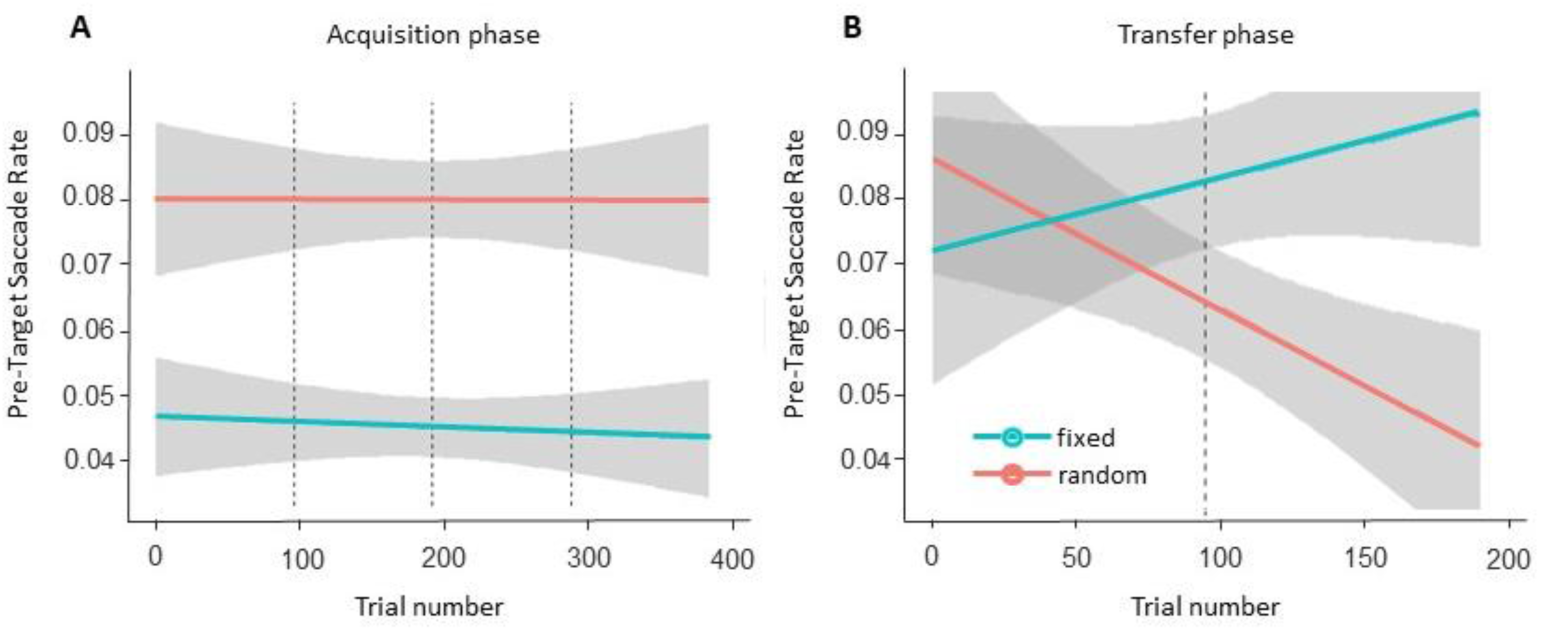
Pre-target saccade rate (SR). Regression line of trial-wise temporal dynamic for pre-target SR in each group (Fixed and Random) during the acquisition (**A**) and the transfer (**B**) phases, based on the individual participant data. Shaded area represents 95% confidence interval around the slope. Dashed lines represent the onset of the different blocks (four acquisition blocks in **A** and two transfer blocks in **B**).

Follow-up analysis within the transfer phase did not reveal a main effect of Group (log estimate= –.404, Wald’s *z*= 1.536, *p*= .125; **Fig 5D**). However, there was a significant interaction between Group and Trial number in this phase (log estimate=. 213, Wald’s *z*= 2.280, *p*= .023): whereas the random groups inhibited their pre-target SR overtime during the transfer phase (log estimate= –.172, Wald’s *z*= –2.327, *p*= .020), the fixed group remained at the same level of pre-target SR over all that phase (log estimate= 0.041, Wald’s *z*= 0.650, *p*= .516; **Fig 4B**). This can be hypothesized to represent the evolvement of an expectation to the new temporal routine over time in the random, but not the fixed group.

**Fig 5.**
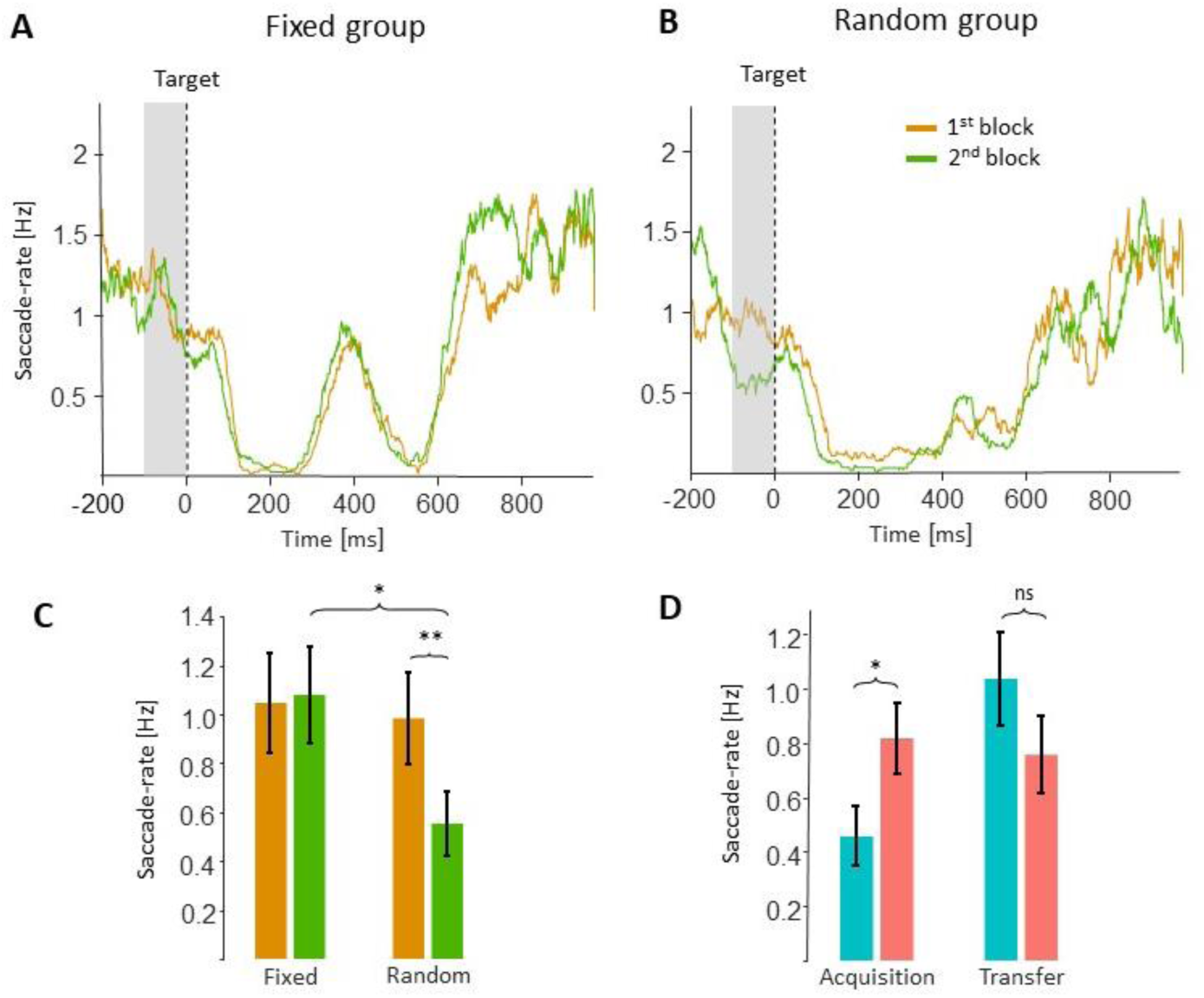
Saccade-rate in the transfer phase. Mean saccade-rate for the first (orange) and second (green) transfer blocks, separately for the fixed (**A**) and the random (**B**) groups. Shadowed area represents the analyzed duration (–100 to 0 ms, pre-target SR), chosen based on previous studies. The dashed lines represent the target onset at time zero. **(C)** Average pre-target SR of the 1^st^ (early) and 2^nd^ (late) blocks, in each group during the transfer phase. *p*<0.05* and *p*<0.01**. **(D)** Average pre-target SR in both phases and groups. Error bars represent ±1 standard error from the group mean. p<0.05*, ns= not significant.

In light of the previous findings (Abeles et al., 2020; Amit et al., 2019; Badde et al., 2020; Dankner et al., 2017; Tal-Perry & Yuval-Greenberg, 2020, 2021) who showed that a decrease in pre-target SR reflects enhanced temporal expectation, the present SR results complement the accuracy-rate findings and suggest that experiencing with fixed vs. random intervals affects the pattern of adjustment to a new regularity.

In addition, a follow-up analysis of the GLMM during the acquisition phase demonstrated a significant difference between groups in this phase (log estimate = –.746, Wald’s *z* = 2.391, *p* = .017), the pre-target SR was higher in the random group compared with the fixed group (**Fig 5D** and **4A**). This effect is consistent with the higher pre-target SR found in the random relative to the fixed condition in previous within-subject studies (Amit et al., 2019; Tal-Perry & Yuval-Greenberg, 2020, 2021) and is a first-time demonstration of this effect between groups.

### Computational modeling: drift-diffusion model

The Drift-Diffusion Model (DDM; Forstmann et al., 2016; Ratcliff & McKoon, 2008; Voss et al., 2013) was used to decompose task performance measures into interpretable psychological processes, in order to better identify the mechanisms underlying the effects obtained for accuracy and RTs. The DDM assumes that evidence is integrated across time until reaching an internal decision-boundary, then a decision is triggered (Glickman et al., 2022; Glickman & Usher, 2019; Zhang, 2012). Here, we compared two variants of the DDM: the first model (hereafter *phase-constant model*) was a simple DDM with three free parameters: i) *Drift-rate* (*v*), indicating the signal-to-noise ratio (SNR) of the evidence, ii) *Boundary-separation* (*a*), indicating the evidence required to make a response, and iii) *Non-decision time* (*t0*), indicating the time of the processes that are not involved in the evidence accumulation (stimulus encoding and response execution). The second model (hereafter *phase-varying model*) was similar to the first model, except that these three parameters were allowed to vary across the acquisition and transfer phases (resulting in six free parameters). Note that, since the drift rate, boundary separation, and non-decision time were estimated separately for the acquisition and transfer phases, the phase-varying model can capture differences in the processing of the sensory stimuli between the acquisition and transfer phases. The models were fitted to the data of each subject using the Fast-DM 30.2 program (Voss & Voss, 2007).

The models were compared using the Bayesian Inference Criterion (BIC), which implements a trade-off between model goodness of fit and complexity by penalizing additional free parameters (Schwarz, 1978). Model comparison showed that the phase-varying model decisively outperformed the phase-constant model (ΔBIC_phase-constant – phase-varying_= 836; values higher than 10 are considered strong evidence), indicating that phase significantly affects the cognitive process underlying the decision. Furthermore, to validate the phase-varying model, we simulated this model using the best-fitted parameters of the participants. The simulated data well captured the decrease in accuracy in the transfer-phase, and in particular the larger decrease shown by the fixed group (**Fig. 6A**). In addition, the phase-varying model well captured the RT patterns of the actual participants in both groups and phases (**Fig. 6B**).

**Fig 6.**
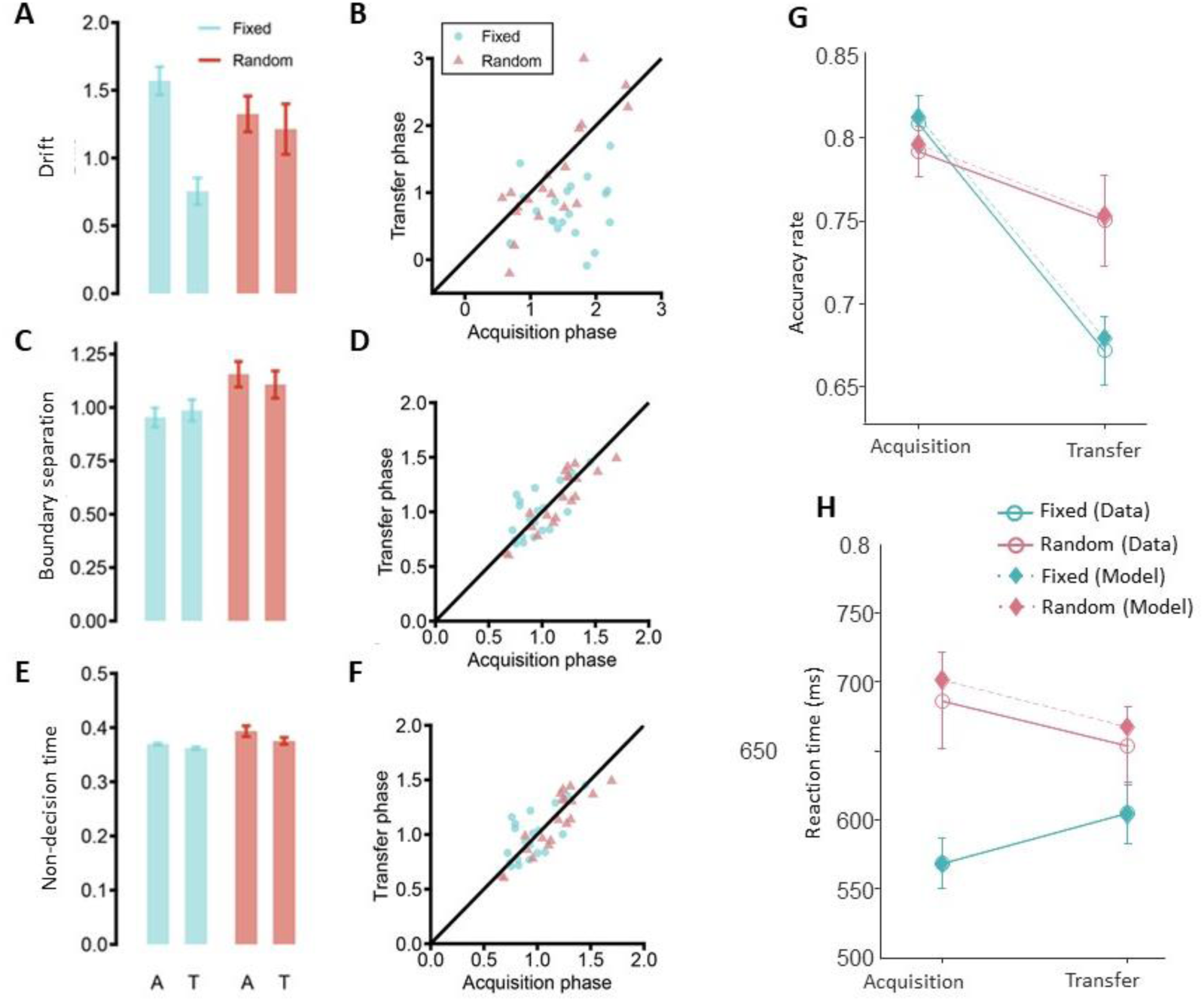
Drift Diffusion Model results. (**A-B**) Accuracy and RTs of the fixed (blue) and random (red) groups in the transfer and acquisition phases for both the real data (solid lines) and model predictions (dashed lines). Error bars represent within-subjects SE. **(C)** Mean drift fit for the fixed (blue) and random (red) groups for the acquisition (A) and transfer (T) phases. Error bars represent ±1 standard deviation from the condition’s mean; **(D)** Single-subject drift fit for the fixed (blue) and random (red) groups for each phase, with the diagonal line representing the identity line (drift fit in acquisition phase = drift fit in transfer phase); **(E-F)** Mean and single-subject boundary separation fits; **(G-H)** Mean and single-subject non-decision point fits. The significant effects between phases within group are marked: p<0.01**, ns= not significant.

Finally, we compared the best-fitted parameters of the phase-varying model obtained in the acquisition and transfer phases, separately for the participants in the fixed and random groups (see Supplementary information S3 for parameter recovery). We found that, for participants of the fixed group, the drift rate (indicating SNR) was significantly higher at the acquisition stage than in the transfer phase (*t*(19) = 5.99, *p* < .001); but no such effect was found for the random group (*t*(18) = .98, *p* = .33). This indicates that the ability of the fixed group to process the target deteriorated once the temporal regularity changed at the onset of the transfer phase, whereas the ability of the random group to process the target remained unchanged. In addition, as a result of the staircase procedure, no difference between the drift rate of the fixed and random groups was found (*t*(37) = 1.47, *p* = .15) in the acquisition phase (indicating a similar SNR). Most importantly, in the transfer phase, the drift rate of the fixed group was significantly lower than that of the random group (*t*(37) = 2.20, *p* = .03), indicating poorer ability to process the sensory input.

No significant differences were found for the boundary parameters between the acquisition and transfer phases for both the fixed (*t*(19) =.83, *p* = .42) and the random (*t*(18) = 1.62, *p* = .12) groups. The non-decision time was higher in the acquisition phase than in the transfer phase, for the fixed group (*t*(19) = 2.72, *p* = .01), and a marginally significant trend in the same direction was found for the random group (*t*(18) = 1.98, *p* = .06). This could possibly result from an improvement in the motor execution due to practice effect.

## Discussion

In this study, we examined how prior experiences with a temporally random environment affects adaptation to a new temporal regularity. Our findings show that prior exposure to a random, relative to a regular temporal environment, enhances adaptation to a new regular environment and promotes the formation of temporal expectations. Participants who were previously exposed to a random temporal pattern adapted more effectively to a new fixed temporal pattern, than participants who were previously exposed to a fixed (but different) temporal pattern. While both groups of participants showed an initial decline in performance following the change in temporal pattern, those who were exposed to the random pattern recovered from this decline and showed an enhancement of performance between the first and the second blocks of the transfer phase, indicating that they have adjusted to the new time routine. In contrast, participants who were exposed to a fixed temporal pattern did not recover from this initial drop. A drift-diffusion model, fitted to the behavioral data (accuracy rates and RTs), suggested that this effect was due to enhancement of sensory processing (i.e., higher SNR level) in the random group relative to the fixed group. This indicates that the implicit learning of temporal regularities is specific to the learned timing routine, and that prior exposure to a random, rather than a fixed, temporal routine enhanced generalizability in a changing environment.

### Eye movements

Previous studies revealed that when a target is predictable, there is a decrease in saccade rate (SR) shortly prior to the predicted target onset (pre-target SR), relative to when the target is unpredictable (Amit et al., 2019; Dankner et al., 2017). This saccade reduction, termed the *pre-target saccadic inhibition,* was found in various, even non-visual modalities (Abeles et al., 2020; Badde et al., 2020) and is thought to reflect anticipation processes (Tal-Perry & Yuval-Greenberg, 2020, 2021). Here, we found that the pre-target SR of the random group decreased between blocks of the transfer phase, and this effect was not observed for the fixed group.

Following the previous studies, we propose that these findings reflect the evolvement of pre-target anticipation in the random group, but not in the fixed group. Consistently with the behavioral results, these findings suggest that prior experience with a fixed regularity hindered participants’ ability to construct new temporal expectations based on a new regularity.

### Learning in contexts of variability

Our findings suggest that prior exposure to temporal uncertainty enhanced adaptation to a new timing regularity. This is consistent with previous studies that demonstrated the importance of noise and uncertainty for learning processes in a variety of human and non-human learning systems. Machine learning studies suggest that adding a certain amount of noise to a trained dataset could enhance the performance of learning algorithms on untrained datasets (Cobbe et al., 2019; Nichol et al., 2018). This effect was offered as a solution to the “over-fitting problem”, by which learning algorithms are trained to fit an existing environment, but fail to generalize when this environment changes (Bishop, 1995a; Reed & Marks, 1999).

Training in variable conditions was found to affect human performance as well. For example, it was found that interleaved practice, also known as *roving*, in which the participants train multiple stimulus types and tasks could hamper immediate learning (Adini et al., 2004) but enhance subsequent generalization (Brady, 1998; Magill & Hall, 1990; Szpiro et al., 2014; Wright et al., 2010). It was further suggested that, under random conditions, participants go through deeper and more elaborate processing of the skill or perceptual task that they practiced (Albaret & Thon, 1998). This hypothesis was supported by fMRI findings showing higher activity in sensorimotor and premotor regions when participants practiced a random variety of motor skills relative to practicing only one skill at a time (Cross et al., 2007). Our present findings are consistent with these studies and extend them to show that an unstable or random environment is beneficial for learning. It should be noted that, unlike most studies of perceptual or procedural learning, in our design, prior experience was very short. Whereas studies of learning typically use repeated-sessions designs, our study was conducted in one session and the acquisition phase included merely four blocks. The effects of long vs. short exposures to randomicity can be examined in future studies.

However, these previous studies on learning in contexts of uncertainty examined the effects of manipulating the level of variability of the trained perceptual feature or task (Greenlee & Magnussen, 1988). In contrast, this study examined the effect of uncertainty in the temporal environment rather than in the trained perceptual features. The oriented stimuli and task demands were constant throughout the experiment and, instead, we manipulated the temporal component of the task, a feature that is only indirectly relevant for performance of the perceptual task. Our findings show that uncertainty of the temporal environment enhanced adaptation to a changing environment. A similar approach was taken by Harris & Sagi (2015) who found that adding temporal variance during training while preserving stimuli features, resulted in better generalization of location.

### A Bayesian interpretation

These findings could also be interpreted via the Bayesian framework, suggesting that our experience of the world is shaped by the integration of bottom-up sensory input (likelihood) and top-down expectations, which are based on prior knowledge on the probabilistic distribution of the target (De Lange et al., 2018; Geisler & Diehl, 2003; Guo et al., 2004; Knill & Richards, 1996). According to this model, the balance between prior and likelihood is based on precision – we prefer to rely on information that is more precise, i.e., has a narrower distribution. Thus, when sensory information is ambiguous, we rely more on our prior knowledge, and when priors are inconsistent, we rely more on sensory evidence (Körding et al., 2004; Yon & Frith, 2021).

In our experiment, the fixed group was presented with a narrow distribution of foreperiods during the acquisition phase, while the random group was presented with a wider distribution. It could be hypothesized that, when integrating sensory information with prior knowledge of the foreperiod distribution, participants of the fixed group relied more on the prior (the acquisition phase distribution) than the random group. During the transfer phase, when encountered with a timing system that was new to both groups, the fixed group faced a large and interfering gap between the new system and their highly-weighted prior. The random group also faced a gap, but it was smaller and less interfering than that of the fixed group. Therefore, during the transfer phase, the random group relied more on current sensory information than on the priors and, consequently, it was easier for them to adjust their predictions according to the new environment. This interpretation is consistent with findings of the drift-diffusion model – participants of the random group perceived the target at a higher SNR level than participants of the fixed group. These findings are consistent with studies who showed that timing behavior was compatible with the prediction of the Bayesian model (Cicchini et al., 2012; Jazayeri & Shadlen, 2010; Miyazaki et al., 2005). This supports the hypothesis that the Bayesian model is fundamental to various aspects of human sensorimotor control and learning (Körding & Wolpert, 2004).

Furthermore, the notion that previous experiences (i.e. priors) are crucial for shaping temporal expectation is consistent with the multiple trace theory of temporal preparation (MTP), as proposed by Los et al. (2014; 2017). According to this theory, the source of temporal expectation are the memory traces that are formed through experience and encode previous timing rules. These memory traces are thought to be aggregated and retrieved from trial to trial (Los et al., 2017). As the Bayesian theory, MTP stresses the importance of prior exposures on the enhancement of performance that is experienced when previously-presented timing regimes are encountered. Consistently with this view, our findings support the involvement of prior experiences and memory trace in the updating of temporal expectations according to Bayesian rules.

## Conclusion

We conclude that, in the case of temporal regularities, learning that the world is systematic could interfere with adaptation to new systems and may be counter-productive for perceptual performance. The finding that training in a temporally-unexpected environment is beneficial, has far-reaching implications for research on the effect of noise in human and computerized learning systems.

## Acknowledgement

This study was funded by ISF grant 1960/19.

## Materials availability statement

The pre-registration file, power analysis report, and data sets are provided at https://osf.io/4g9px/. For the statistical and experimental codes, please contact.

## Methods

### Participants

Participants included 40 students. Participants were Cucasians of Arab and Jewish decent, corresponding with the ethnic characteristics of most student population of Tel Aviv University. The participants were randomly divided into two groups of 20 participants each (Random group: 12 females, age: Mean 24.73, SD 3.71, Range 18-33; Fixed group: 11 females, age: Mean 25.7, SD 5.25, Range 19-43). One of the participants (random group) failed to perform the task during the acquisition phase, as indicated by mean accuracy of 58.85% during that phase, and was therefore excluded from analysis, based on the pre-registered exclusion criteria (link: https://osf.io/td5z3).

The sample size was determined based on a power analysis simulation of pilot data of three participants in the fixed group and four participants in the random group. These data were not included in the main study (as mentioned in the preregistration document; Power = 0.80, α = 0.05). All participants reported normal or corrected-to-normal vision and no history of neurological disorders. All were naïve to the purpose of this study. They participated in the experiment in exchange for course credit or monetary compensation. The ethical committees of Tel Aviv University and the School of Psychological Sciences approved the study. Before participation, all participants signed informed consent.

### Stimuli

The stimuli included a black or blue fixation cross (0.08 x 0.08°) on a mid-gray background. An auditory cue consisted of a pure tone of 5 KHz, played for 25 ms. The target was a slightly tilted Gabor grating patch (visible diameter 3°, spatial frequency 5 cycles per degree, contrast 0.7). The tilt degree was determined individually using an adaptive staircase procedure. The target stimulus was followed by a mask, composed of two overlaid Gabor grating patches (same parameters as the targets) in full contrast, tilted at 45 and 315 degrees.

### Procedure

*Setup*. Participants were seated in a dimly lit sound-attenuated chamber, head resting on a chinrest at a distance of 100 cm from the display monitor (24” LCD ASUS VG248QE, 1,920 × 1,080 pixels resolution, 120 Hz refresh rate, mid-gray luminance was measured to be 110 cd/m^2^). Auditory stimuli were administered through speakers located on both sides of the monitor. The experiment was generated and controlled using MATLAB R2015a (MathWorks Inc., Natick, MA, USA) with Psychophysics Toolbox v3 (Brainard, 1997).

*Trial Procedure.* Each trial began with the presentation of a central black fixation cross. Following 0.5 s of stable fixation (<3° off center) as verified by a gaze contingent procedure and a random inter-trial interval (ITI) of 0.2-0.7 s (uniformly distributed), the fixation cross changed color to blue and, simultaneously, a pure tone was heard. Both the visual and the auditory signals served as concurrent temporal cues marking the onset of a foreperiod. The blue fixation cross remained on the screen throughout the foreperiod interval, and the auditory tone was paused after 25 ms. The reason for using both visual and auditory signals as cues was to increase the strength of the temporal information and thus enhance the formation of temporal expectation for the target stimuli (Menceloglu et al., 2017).

The foreperiod lasted for varying durations, depending on the group and the experimental phase (see below). Following the foreperiod, the target stimulus was briefly presented (for 50 ms) at the center of the screen and after 50 ms of a blank screen, the backward mask was presented centrally for 300 ms. Participants were asked to report the orientation of the target’s tilt (clockwise / counterclockwise) by pressing one of two keys. When a response was detected or once 3 s had elapsed with no response, the fixation changed color again to green or red for 500 ms to provide feedback on task performance (correct or incorrect, respectively). A lack of response was considered as an incorrect response. Following the feedback, the next trial was initiated. The trial procedure is depicted in **Fig. 1**.

*Staircase procedure.* A 1-up 3-down staircase procedure was performed on the tilt of the target stimulus during a separate session before the main experiment to determine individual thresholds. The purpose of the staircase session was to ensure that the task difficulty level of both groups would be similar and to avoid ceiling or floor effects. The starting tilt of the Gabor-patch was set at 10° and participants performed three staircase blocks of five minutes each, using a staircase procedure, with logarithmic steps, which yielded accuracy rates of approximately 80%. During the staircase procedure, foreperiods distribution was identical to the participant’s acquisition phase, determined by their group. The average tilt of the last eight reversals was taken as the participant’s tilt for the experimental blocks (both for the acquisition and the transfer phases). Following the pre-registered exclusion criteria, participants who failed to perform the task during the staircase procedure (performance at the chance level) or were assigned a tilt of more than 10° following the staircase procedure were excluded from the experiment and did not proceed to the main experimental session. The resulting average tilt of the two groups was nearly identical (*fixed group*: 3.283°±2.252° SD; *random group*: 3.135°±1.525°). Twenty-six of the participants (divided equally between groups) did not perform any training in addition to the staircase procedure.

However, while running the experiment we encountered some participants who were unable to perform the task, and therefore, we added 12 training trials before the staircase procedure, for the rest of the participants (14 participants). Two of the training trials were without a mask, to ensure that participants were familiar with the target and understood the task. This short training did not affect the tilt threshold or the performance during the acquisition or transfer phases. The full analysis of the staircase procedure can be found in Supplementary materials S1.

*Experimental blocks.* The main experimental session began with four blocks of the acquisition phase, followed by two blocks of the transfer phase (see **Fig 1B**), with each block consisting of 96 trials and breaks administered between them. During the *acquisition phase*, the foreperiod varied by group and was either constant at 2.7 s for the *fixed group* or sampled randomly from a continuous uniform distribution ranging between 1.7 and 3.7 s (mean 2.7 s) for the *random group*. During the *transfer phase*, foreperiod distribution was identical for both groups and included 80% of the trials with foreperiod of 0.7 s, and 20% of the trials with foreperiod of 2.7 s (in random order). Instructions were provided before the first block. The accuracy rate of the previous block was presented during each break to facilitate task engagement.

### Eye tracking

Binocular gaze position was monitored using a remote infrared video-oculographic system (Eyelink 1000 Plus; SR Research, Canada), with a spatial resolution ≤0.01° and average accuracy of 0.25°–0.5° when using a headrest (as reported by the manufacturer). Raw gaze positions were sampled at 1000 Hz and converted into degrees of visual angle using a 9-point-grid calibration, which was performed at the beginning of the experimental session. Saccades were detected using a modification of a published algorithm (Engbert, 2006), which was applied on filtered gaze position data (low-pass IIR Butterworth filter; cutoff 60 Hz; as in Amit et al., 2017). An elliptic threshold criterion for microsaccade detection was determined in 2D velocity space based on the horizontal and vertical velocities of eye-movement. We segmented the recorded eye movement data and created a segment of 2000 ms per trial starting from –1000 ms and ending at +1000 ms relative to target onset at time zero. The median SD of the eye movements velocity was estimated based on these segments, and the microsaccade detection threshold was set to be six times the SD. Saccade onset was defined when six or more consecutive velocity samples were outside the ellipse, in both eyes. Saccades offsets are sometimes accompanied by an overshoot, which may be erroneously detected as a new saccade. Therefore, per standard procedure, we imposed a minimum criterion of 50 ms for the interval between two consecutive saccades and kept only the first saccade in cases where two saccades were detected within such interval. Saccades of all sizes were included in the analysis, but due to the instruction to keep sustained fixation, most saccades were small (in the range of microsaccades <1°).

The time series of saccade rate was constructed as in previous studies (Abeles et al., 2020; Amit et al., 2019; Dankner et al., 2017; Tal-Perry & Yuval-Greenberg, 2020, 2021) for each participant by counting the number of saccade events in each time-point across trials, separately for each group and block, and dividing these values by the number of trials (discounting trials in which a blink was detected in the given time sample).

### Statistical Analysis

*Behavioral analysis.* Accuracy-rates and mean reaction time (RT) were calculated separately for each participant and group. Only correct trials were included in the RT analysis. Trials deviating by more than 3 SD from the mean RT of each participant were also excluded from the data analysis. Analysis of the transfer phase included only the standard trials of foreperiod 0.7 s and not the rare trials of foreperiod 2.7 s.

Accuracy and RT were analyzed using generalized linear mixed models (GLMM) with fixed factors of Group (fixed/ random), Phase (acquisition /transfer), and the phase’s trial ID (continuous). We constructed the GLMM assuming a binomial family of response and logit link (i.e., a logistic mixed effect model) for the accuracy model (Jaeger, 2008) and a Gamma family of response with an identity link for the RT model (Lo & Andrews, 2015).

To capture the temporal dynamics across the phase, the trial ID was modeled using polynomials with increasing polynomial degrees fitted to the data. The polynomial models were iteratively compared using likelihood-ratio tests until there was no significant improvement in fit (*p* > .05). Treatment contrasts were used for both Group and Phase factors, setting the fixed group and the acquisition phase as the base levels. Fixed effects were tested by comparing the effect against a reduced model without the effect (akin to type-II sum-of-squares, SS), using likelihood-ratio tests. Simple effects were examined using parameter coefficients and tested using the Wald z-test and the base levels were reset to examine some of the models’ simple effects. To balance type-I and type-II errors (Matuschek et al., 2017), the random effect structure of the model was chosen according to the model most parsimonious with the data (Bates et al., 2015a), as follows. First, a model allowing for a random intercept by subject was considered. Next, we tested whether a model with added random slopes by subject for fixed factors provided a better fit. Lastly, we tested whether a model with both random slopes by subject and random interaction slopes by subject better fits the data. In each iteration, we tested, using a likelihood-ratio test (against *a* = .05), whether the model converged, and, additional, whether there was a significant improvement in model fit relative to the reduced model. Models that failed to converge were trimmed from their random slopes or random interaction slopes, beginning with the random term that explains the least amount of variance, until reaching convergence.

*Eye-movements analysis*. In a series of previous studies, it was established that pre-target inhibition of saccade-rate reflects temporal expectation to a predicable target. Specifically, saccades were found to be inhibited shortly before predictable, but not unpredictable, targets (Abeles et al., 2020; Amit et al., 2019; Badde et al., 2020; Dankner et al., 2017; Tal-Perry & Yuval-Greenberg, 2020, 2021). Here, we used this index of pre-target saccade rate (SR) to examine the hypothesis that participants of the random group would show better adjustment, i.e., higher temporal expectation, to the new regularity during the transfer phase. To this end, we examined the mean pre-target SR: mean saccade rate at –100 to 0 ms relative to target onset, as was done in previous studies and was pre-registered (Abeles et al., 2020). Since this measurement is affected by the presentation of the cue more in trials with short foreperiods than in trials with long foreperiods, we focused this anlaysis on comparing the same foreperiod between groups.

In order to examine changes in SR across time, we calculated the mean pre-target SR (at –100 to 0 ms) using a generalized linear mixed model (GLMM) for the pre-target SR. This model was constructed similarly to the accuracy and RT GLM models (see above). Another, subsidiary, goal of this analysis was to test whether the finding of enhanced SR inhibition for predictable, relative to unpredictable, targets holds even in a between-subject design. Since one group was presented with random foreperiods and the other with fixed foreperiods in the acquisition phase, this goal was achieved by comparing the pre-target SR of the two groups, in this phase.

*Analysis software.* GLMM analyses were done in R v4.0.3 using R-studio v1.3.959. We used the lme4 (Bates et al., 2015b) package in R, with model diagnostics done with the performance package (Lüdecke et al., 2020). The behavioral outcomes and eye-movements results were processed in MATLAB R2020a (The MathWorks Inc., Natick, MA, USA). The statistical analyses were done in JASP v0.14 (JASP Team, 2020).

## Notes

https://osf.io/4g9px/

### Competing Interest Statement

The authors have declared no competing interest.

