## supplementary material for "Exposure to temporal randomness promotes subsequent adaptation to new temporal regularities"

### Supplementary materials

***Supplementary material S1: Staircase procedure analyses***

Prior to the main experiment, participants performed a staircase procedure aiming to determine the tilt threshold for each participant individually and to detect participants who failed to perform the task and should be excluded from the study. To confirm there was no difference in staircase results between groups (fixed/ random), the threshold Gabor tilt angles were compared using an independent *t*-test, and no evidence for a difference between groups was found ($t(38) = -.003$, $p = .997$).

In addition, we examined whether additional training performed by some of the participants affected the results. After the first participants experienced difficulties in task performance, we added twelve additional trials of training prior to the staircase procedure for the remaining participants. To ensure that this additional training did not change the results, we compared participants who performed this extra training (N=26), with those who have not (N=14). We found no difference for threshold tilt angles between these two groups ($t(37) = 1.349$, $p = .185$), and no interaction between training condition and group (fixed vs. random; $F(1) = 1.731$, $p = .197$). In addition, we found no evidence for differences in performances between them during the experimental phases (*Acquisition phase*: Accuracy-rates: $t(37)$ = .736, $p$ = .467;, RT: $t(37)$ = .022, $p$ = .983; *Transfer phase:* Accuracy-rates: $t(37)$ = .200, $p$ = .843, $t(37)$ = -.462, $p$ = .647).

***Supplementary material S2: Analyses that were declared in the pre-registration document but were not included in the main manuscript***

*Eye-movements.* Besides the GLMM analysis described in the main text, we employed a pre-registered 2X2 mixed ANOVA with a between-subject factor of Group (fixed /random) and a within-subject factor of Block (first/ second) on the pre-target SR in the transfer phase. The results of the GLMM for pre-target SR (see **Results**) are consistent with the findings that are described below.

We found no significant main effect for Group ($F(1, 35)$= 1.543, $p$=.222) nor for Trial bin ($F(1, 35)$= 3.366, $p$=.075), but there was a significant interaction between them ($F(1, 35)$= 4.580, $p$=.039), indicating that pre-target SR decreased over time in the random group, while it remained constant or even slightly increased in the fixed group (**Fig 4B and 5D)**. Follow-up analysis demonstrated that at the beginning of the transfer phase (the first block), there was no significant difference between the groups ($t(35)$=0.219, $p$=.828), but at the end of the transfer phase (the second block), pre-target SR was higher for the fixed relative to the random groups ($t(35)$=2.202, $p$=.034). When examining the effect of time on the two groups separately, we found that the pre-target SR of the random group decreased dramatically between the first and the second transfer blocks ($t(18)$=3.385, $p$=.003), whereas that of the fixed group has not changed significantly ($t(19)$=0.192, $p$=.850; **Fig 5C**).

In addition, we compared the pre-target SR of the groups during the acquisition phase using an independent samples t-test and found a significant difference between the groups ($t(37)= -2.081$, $p=.044$; **Fig. 5D and 4A**): pre-target SR was higher for the random relative to the fixed group. This effect is consistent with the higher pre-target SR found in a random relative to a fixed condition in previous within-subject studies (Amit et al., 2019; Tal-Perry & Yuval-Greenberg, 2020, 2021) and is a first-time demonstration of this effect between groups.

*Reaction times.* Our design was not suitable for studying RT, because we used a difficult, not speeded task. However, to follow the pre-registered plan, and to provide further evidence for the lack of speed-accuracy tradeoffs, we report here the full RT results.

RTs were examined similarly to the accuracy-rates, using a three-way generalized-linear-mixed-model (GLMM) but assuming a Gamma response family with an identity link. The same fixed factors as in the accuracy analysis were used, except that the Trial number dynamics were best fitted using $1^{st}$ and $2^{nd}$ polynomials. The model allowed for a random intercept and random slope for phase by subject.

We found no main effect for Phase ($\chi^2(1) = .008$, $p = .927$) and only a marginal effect for Group ($\chi^2(1) = 2.929$, $p = .087$), but the main effect of Trial number was significant ($\chi^2(2) = 56.298$, $p < .001$), as participants became faster overtime. As in the accuracy-rates findings, there was a significant three-way interaction ($\chi^2(2) = 16.597$, $p < .001$) of Phase, Group and Trial number. Also, all the two-way interactions were found to be significant (Trial number X Group: $\chi^2(2) = 22.015$, $p < .001$; Group X Phase: $\chi^2(1) = 4.389$, $p = .036$; Trial number X Phase: $\chi^2(1)$

= 75.881, $p < .001$). A follow-up examination revealed that during the acquisition phase, the groups showed different trends of RT overtime. The fixed group became faster (log estimate = -8.589, Wald's $z = -7.277$, $p < .001$) while the random group became slower (log estimate = 6.627, Wald's $z = 5.142$, $p < .001$). These different trends of RT cannot explain the accuracy findings, as accuracy rates were similar between groups during that phase (as a result of the staircase procedure). During the transfer phase both groups presented similar and significant trends of decreasing RT overtime (random group: log estimate = -16.542, Wald's $z = -11.163$, $p <.001$; fixed group: log estimate = -17.910, Wald's $z = -15.663$, $p<.001$; **Fig 3**), providing no support for a speed-accuracy tradeoff.

*Dependency of the effect on performance in the acquisition phase.* In the preregistration document, we suggested to perform a complementary analysis that examine the effect of task performance during the acquisition phase on accuracy rates of the transfer phase. The purpose of this analysis was to test the hypothesis that an effective prior exposure, manifested by high performance during the acquisition phase, would enhance adjustment to the new regularity of the transfer phase, relative to less effective exposure. To examine this hypothesis, each of the two groups was split into two sub-groups ('good' performers and 'poor' performers), based on median split calculated separately for each group on the average accuracy rates at the acquisition phase. A three-way mixed ANOVA with between subject factors of Acquisition-group (fixed/ random) and Quality-group (good/ poor) and a within subject factor of Block (transfer block T1/ T2) was performed on the accuracy rates. Following the aim of this analysis, we focused only on interactions with the factor Quality-group.

There were no significant interactions of any of the factors with the factor Quality-group (Block X Quality-group: $F(1, 35) = .013$, $p = .910$; Quality-group X Acquisition-group: $F(1, 35) = 1.779$, $p = .191$; Three-way interaction: $F(1, 35) = 1.015$, $p = .321$). Yet, when exploring the 'good' and 'poor' performers separately (**Fig. S1**), we found that the difference in performance between the random and fixed groups during the transfer phase was evident only for the good performers ($F(1) = 7.635$, $p = .013$) and not for poor ones ($F(1) = .379$, $p = .546$). This indicates that a minimal level of engagement and task compliance is necessary in order to achieve the observed modulation at the transfer phase. This finding further strengthens the link between experience during the acquisition phase and performance during the transfer phase.

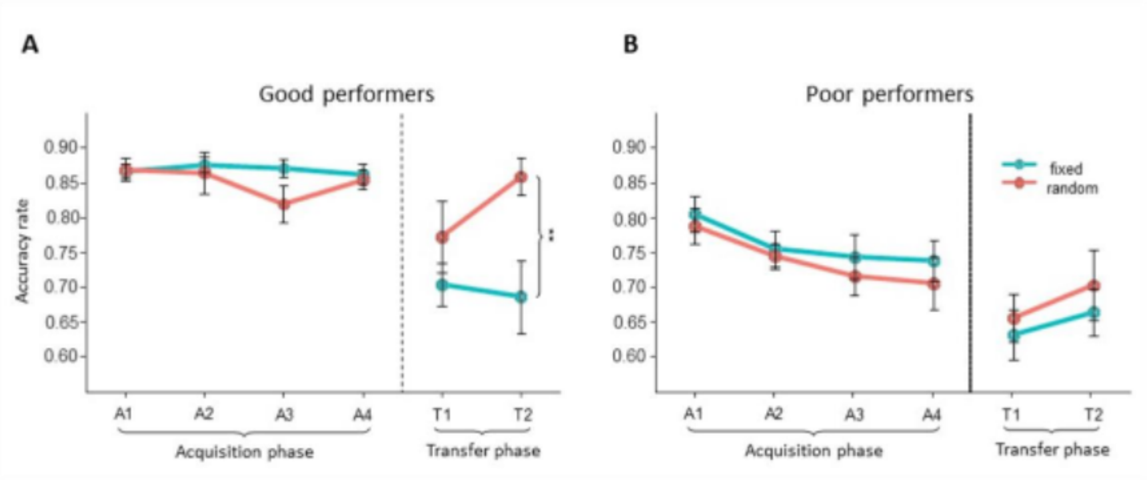


**Fig. S1: Accuracy rate for good (A) and poor (B) performers**. The X-axis represents the experimental blocks in chronological order (Acquisition= A 1-4, Transfer= T 1-2). Error bars depict ±1 standard error from the group mean. Simple effects are marked: p<0.01**. Dashed line represents the onset of the transfer phase

### ***Supplementary material S3: Drift diffusion model- Parameter recovery***

Parameter recovery was performed for the phase-varying drift-diffusion model (our best-fitting model). The model was first simulated using the estimated parameters of each participant (generative parameters) and the same number of trials as there is in the data. Then, the model was fitted to the simulated data and its parameters were extracted (recovered parameters). **Fig. S2** shows the generative parameters plotted against the recovered parameters. As shown, the quality of the parameter recovery can be considered as excellent (White et al., 2018).

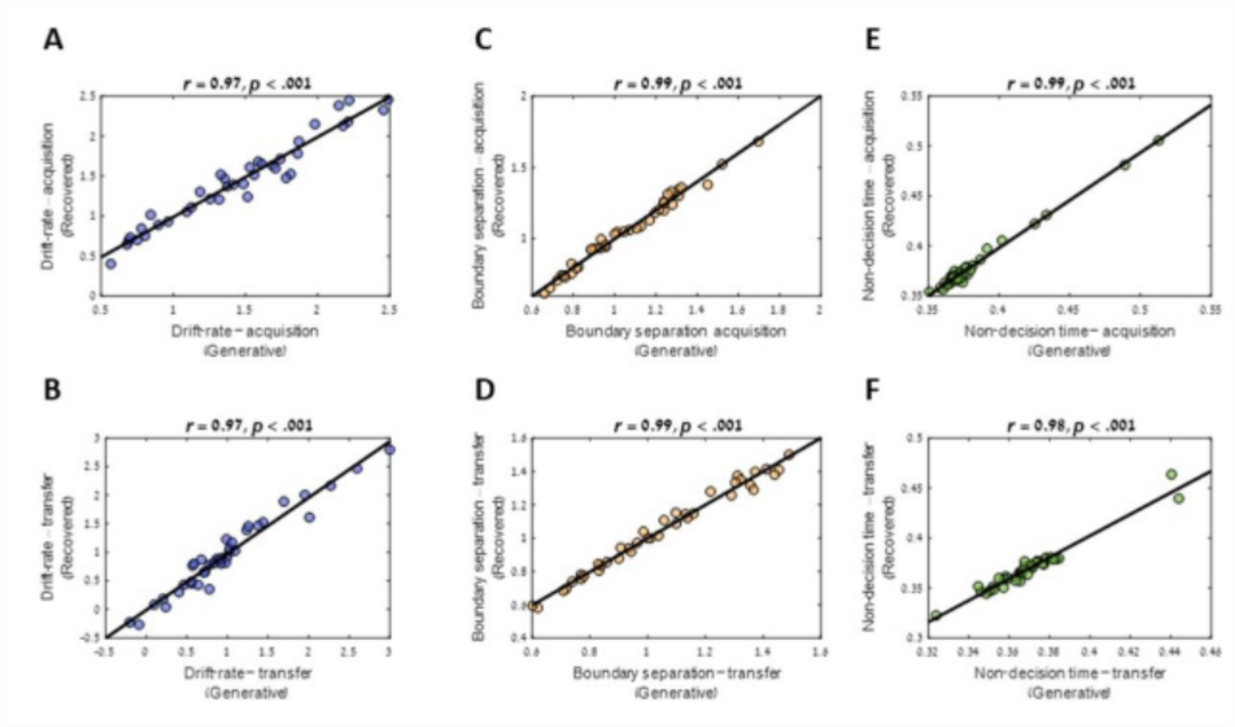


**Fig. S2. Parameter recovery for the phase-varying drift-diffusion model.** The generative (A) drift-rate – acquisition phase, (B) drift-rate – transfer phase, (C) boundary separation – acquisition phase, (D) boundary separation – transfer phase, (E) non-decision time – acquisition phase and (F) non-decision time – transfer phase are plotted against the recovered corresponding parameters.
